# Large-scale monitoring of resistance to coumaphos, amitraz and pyrethroids in *Varroa destructor*

**DOI:** 10.1101/2020.11.12.378190

**Authors:** Carmen Sara Hernández-Rodríguez, Óscar Marín, Fernando Calatayud, María José Mahiques, Ana Mompó, Inmaculada Segura, Enrique Simó, Joel González-Cabrera

**Affiliations:** Instituto Universitario de Biotecnología y Biomedicina (BIOTECMED), Universitat de València. Dr. Moliner 50. 46100 Burjassot, Spain; Agrupación de Defensa Sanitaria Apícola APIADS. Calle Raval 75B. 46193 Montroi, Spain; Agrupación de Defensa Sanitaria Apícola APICAL y APIVAL. C/Sants de la Pedra 75. 03830 Muro de Alcoy, Spain

**Keywords:** acaricides, honey bees, bioassay, TaqMan, genotyping, acaricides

## Abstract

*Varroa destructor* is an ectoparasitic mite causing devastating damages to honey bee colonies around the world. Its impact is considered a major factor contributing to the significant seasonal losses of colonies recorded every year. Beekeepers are usually relying on a reduced set of acaricides to manage the parasite, usually the pyrethroids tau-fluvalinate or flumethrin, the organophosphate coumaphos and the formamidine amitraz. However, the evolution of resistance in the populations is leading to an unsustainable scenario with almost no alternatives to reach an adequate control of the mite.

Here we present the results from the first, large-scale and extensive monitoring of the susceptibility to acaricides in the Comunitat Valenciana, one of the most prominent apicultural regions in Spain. Our ultimate goal was to provide beekeepers with timely information to help them decide what would be the best alternative for a long-term control of the mites in their apiaries. Our data show that there is a significant variation in the expected efficacy of coumaphos and pyrethroids across the region, indicating the presence of a different ratio of resistant individuals to these acaricides in each population. On the other hand, the expected efficacy of amitraz was more consistent, although slightly below the expected efficacy according to the label.

**HIGHLIGHTS:**

1. *Varroa destructor* is causing severe damages to honey bee colonies worldwide.
2. There are very few acaricides available to manage the parasite.
3. The evolution of resistance is limiting our capacity to control the mite.
4. We estimated the expected efficacy of the main acaricides in many Spanish apiaries.
5. The information was shared with beekeepers for them to decide the best treatment to control the mite.

## INTRODUCTION

The ectoparasitic mite *Varroa destructor* is considered a major pest of the Western honey bee (*A. mellifera* L.) (Rosenkranz et al. 2010). This mite feeds mostly on the fat body of immature and adult bees and vectors numerous lethal viruses (Boecking and Genersch 2008, Ramsey et al. 2019), compromising the natural honey bee defences. These severe damages make *V. destructor* one of the main factors contributing to the many seasonal losses of honey bee colonies around the world (Genersch et al. 2010, Guzmán-Novoa et al. 2010).

*Varroa destructor* shifted host from the Eastern honey bee (*Apis cerana* L.) to the Western honey bee in the late 1950’s in Asia, but nowadays it is widely distributed throughout the world (Solignac et al. 2005). In Spain, *V. destructor* was first detected in 1985 and currently it can be found all over the country (Llorente 2003, Muñoz et al. 2015).

*Varroa destructor* reproduces throughout the spring and summer, so the population is larger in autumn. Thus, treatments to control de mite are usually applied in that season to increase the possibility of overwintering success (Underwood and López-Uribe 2020). In this country, as in many others, it is mandatory to apply at least one acaricide treatment per year to manage the parasite (Royal Decree RD608/2006). However, beekeepers usually perform at least another treatment in summer in case they detect mites in their hives. The acaricides authorised to control *V. destructor* in Spain include “hard acaricides” (based on pyrethroids like tau-fluvalinate and flumethrin; the formamidine amitraz, and the organophosphate coumaphos), together with “soft acaricides” (mostly based on formic or oxalic acids, and the essential oil thymol) (FAO, 2000; www.aemps.gob.es/). Integrated Pest Management (IPM) strategies encourage the combined use of both types of acaricides and other beekeeping practices to reach better long-term control of the mite, but beekeepers are relying mainly on hard acaricides because they are faster and usually more effective (Rosenkranz et al. 2010).

The intensive use of pyrethroids to control Varroa for decades resulted in the emergence of resistance to these acaricides in apiaries from several countries (Milani 1995, Elzen et al. 1998, Sammataro et al. 2005, Gracia-Salinas et al. 2006, González-Cabrera et al. 2018). Since the emergence of *V. destructor* resistance to pyrethroids, beekeepers switched to coumaphos as the best alternative to control the parasite, but the intensive treatment regime with this compound resulted in the evolution of resistance in many locations too (Elzen and Westervelt 2002, Maggi et al. 2009, Maggi et al. 2011). In this scenario, the alternatives to control *V. destructor* have been drastically reduced to the use of amitraz and soft acaricide treatments. Currently, the extensive use of amitraz is exerting an intense selection pressure over populations, threatening them with the evolution of resistance to this compound. Indeed, a reduction in the efficacy of amitraz for Varroa control, probably associated with the evolution of resistance, has already been reported elsewhere (Rodríguez-Dehaibes et al. 2005, Maggi et al. 2010, Kamler et al. 2016, Rinkevich 2020).

The mechanism of resistance to pyrethroids in *V. destructor* is well known. It is associated with mutations at the residue L925 of the major target site for pyrethroids: the Voltage-Gated Sodium Channel (VGSC) (González-Cabrera et al. 2013, Hubert et al. 2014, González-Cabrera et al. 2016, González-Cabrera et al. 2018). To detect these amino acid substitutions, TaqMan® allelic discrimination assays have been developed (González-Cabrera et al. 2013, González-Cabrera et al. 2016, González-Cabrera et al. 2018). This is a high throughput diagnostics technique capable of detecting the mutation in individual mites. On the other hand, the molecular mechanisms causing the resistance to coumaphos and amitraz in *V. destructor* are still unknown, so the reduction in the efficacy to control *V. destructor* in apiaries using treatments based on these two acaricides can only be confirmed by bioassays with a direct exposition of mites to the acaricidal products (Elzen and Westervelt 2002, Maggi et al. 2009, Maggi et al. 2011).

The European Union (EU) is the world’s second largest honey producer after China. Spain is the country with the most hives in Europe (more than three million hives) and the second largest producer of honey in the EU, with almost 30,000 tons per year (https://ec.europa.eu/info/food-farming-fisheries/animals-and-animal-products/animal-products/honey_en). The Comunitat Valenciana is a Spanish extensive region of 23,255 km^2^ comprising three provinces with an important professionalized beekeeping sector. It has a census of 358,327 hives in 2,459 beekeeping operations, being the second region with the highest honey production in Spain. Almost all the hives (98 %) are mobile, carrying out migratory beekeeping throughout the year (https://www.mapa.gob.es/es/ganaderia/temas/produccion-y-mercados-ganaderos/indicadoreseconomicossectordelamiel2018comentarios_tcm30-419675.pdf).

Despite the treatments to manage the mite, Varroa parasitism, far from being controlled, it is a persistent problem in all honey-producing countries. It seems plausible that the continuous presence of varroosis in beehives around the world is related to resistance to acaricidal products. In Spain, this correlation has not been confirmed yet since there is no program to track the efficacy of treatments or the evolution of resistance in Spanish apiaries. The lack of knowledge of the incidence and prevalence of varroosis in all the territories of this country led us to plan a systematic study in the Comunitat Valenciana region. The goal of this study was to determine the efficacy of the three groups of hard acaricides in *V. destructor* populations from apiaries located throughout the three provinces of the region. This study was coordinated with the Department of Agriculture, Environment, Climate Change and Rural Development of the regional government of the Comunitat Valenciana (Generalitat Valenciana, www.gva.es) and Sanitary Defence Groups (ADS acronym in Spanish) of the beekeeping sector. Our aim was to provide beekeepers with information about the impact of varroosis in their colonies and to estimate the possible efficacy for each apiary of acaricide treatments based on pyrethroids, coumaphos and amitraz.

## MATERIAL AND METHODS

### Mites

*Varroa destructor* females were collected from apiaries located in the three provinces of the Comunitat Valenciana region (Spain): Castellón, Valencia and Alicante. The sample collection was carried out in two consecutive annual beekeeping seasons: first, 90 samples were collected from April to July 2018; and in the second period, 82 samples were collected from November 2018 to July 2019. Beekeepers and veterinaries from nine Sanitary Defence Groups (ADS), participated in the collection and shipments of the samples (Table 1). At least two combs with capped brood per apiary were collected, boxed in polystyrene containers and shipped to the laboratory at the University of Valencia by express courier. The samples arrived in less than 24 hours after collection to ensure optimal mite conditions before bioassays.

**Table 1.** Sanitary Defence Groups (ADS) that supplied samples of *Varroa destructor*.

| Sanitary Defence Groups<br>(ADS) | Province | No. hives<sup>1</sup> | No. samples<sup>2</sup> |
| --- | --- | --- | --- |
| ADSAD | Valencia | 16901 | 9 |
| AIXAM | Castellón | 16168 | 9 |
| ALAPI | Alicante | 4957 | 3 |
| APAC | Castellón | 46317 | 25 |
| APIADS | Valencia | 93548 | 50 |
| APICAL | Alicante | 44210 | 27 |
| APIVAL | Valencia | 47332 | 22 |
| CASAPI | Castellón | 14904 | 9 |
| PROAPI | Valencia | 43656 | 18 |
| TOTAL |  |  | 172 |

<sup>1</sup>Total number of hives belonging to the ADS.
<sup>2</sup>Number of samples provided for this study.

### Bioassays with acaricides

Bioassays were conducted as in Higes et al. (2020) using strips of Checkmite+ (coumaphos a.i., Bayer, Germany), Apitraz (amitraz a.i., Laboratorios Calier, S.A., Spain), Amicel Varroa (amitraz a.i., Maymó S.L., Spain), and Apivar (amitraz a.i., VétoPharma, France). Briefly, parasitized bee pupae were extracted from the brood cells using a pair of soft tweezers (Fig. 1A). The female mites were collected with a soft paint brush and deposited onto a wet filter paper. A piece of approximately 4 cm long of each acaricide strip was placed into a 5.5 cm Petri dish. Amicel Varroa strips were prepared following the manufacturer instructions. Each strip piece maintained its original width (2.5 cm for Checkmite+; 4.0 cm for Apitraz and Apivar; and 4.6 cm for Amicel Varroa). Given the different amount of active ingredient impregnated in the strips of each product, and considering the surface of both sides of the strips, the actual concentration was 13.6 mg/cm^2^, 2.1 mg/cm^2^, 0.8 mg/cm^2^ and 3.1 mg/cm^2^ for Checkmite+, Apitraz, Amicel and Apivar, respectively. The mites collected (15 mites per replicate, 2-3 replicates for each acaricide product) were laid on top of the strip and their movements were monitored to control that they remain on top of the strip for at least 5 min (Fig. 1B). The dish was sealed with Parafilm® and holed with an entomological needle to allow aeration. Mites onto the acaricide strip were incubated for 1 h at 34 °C, 90 % RH in a wet incubator. After 1 h, the strip was removed and the dish with the mites was incubated for 3 more hours at 34 °C, 90 % RH. Controls mites were treated the same way but without acaricide strips. After the incubation time was completed, mortality was evaluated by assessing the movement of mites after probing them with a fine paint brush (Fig. 1C). The expected efficacy of each acaricide was estimated using the mortality values obtained in the bioassays.

**Figure 1.**
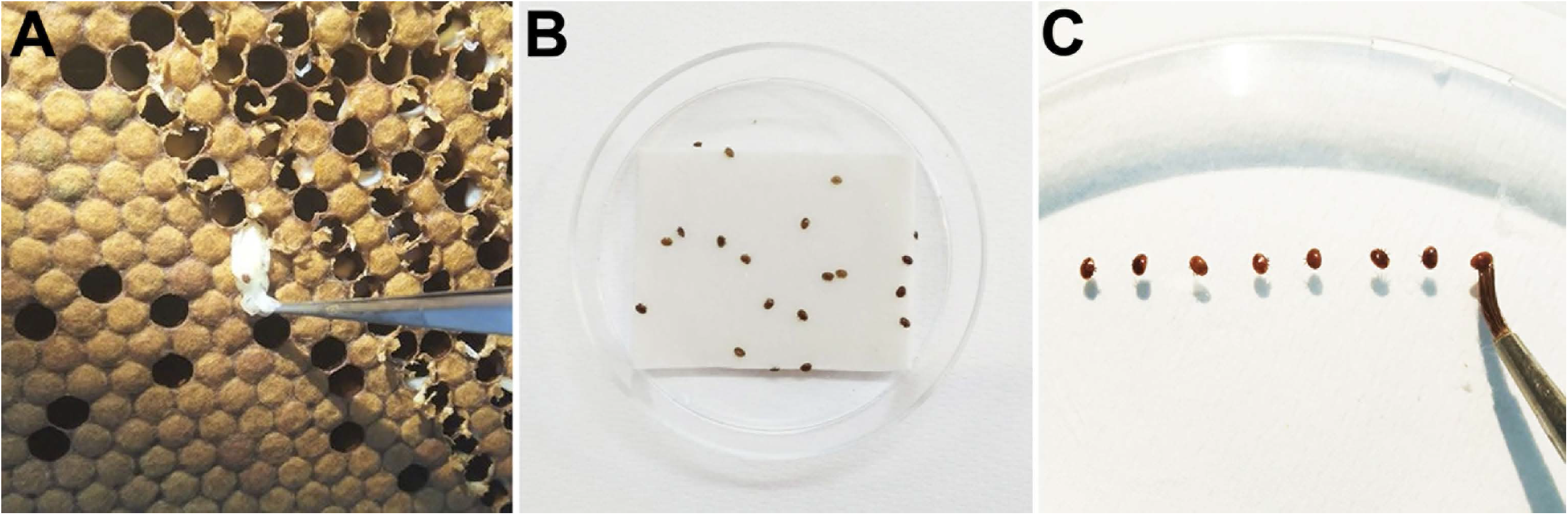
Bioassays with *Varroa destructor*. (A) Parasitized bee nymphs extracted from the brood cells. (B) Female mites laid on the acaricide strip into a Petri dish. (C) Mortality was evaluated by assessing the movement of mites after probing them with a fine paint brush.

### TaqMan® assays

Genotyping of mites for detecting susceptible and pyrethroid-resistant alleles in *V. destructor* VGSC was carried out using a TaqMan® based allelic discrimination assay as described by González-Cabrera et al. (2013). Genomic DNA was extracted from individual adult mites by an alkaline hydrolysis method and stored at −20 °C until used. Briefly, reactions mixtures contained 1.5 μl of genomic DNA, 7.5 μl of TaqMan® Fast Advanced Master Mix (Thermo Fisher Scientific), 0.9 μM of each primer and 0.2 μM of each fluorescent-labelled probe in a total reaction volume of 15 μl. Assays were run on a StepOne Plus Real-Time PCR system (Thermo Fisher Scientific) using the following temperature cycling conditions: 10 min at 95 °C followed by 40 cycles of 95 °C for 15 s and 60 °C for 45 s. The increase in VIC® and 6FAM™ fluorescence was monitored in real time by acquiring each cycle on the yellow channel (530 nm excitation and 555 nm emission) and green channel (470 nm excitation and 510 emission) of the StepOne Plus, respectively. Forty mites were genotyped per apiary. Duplicated control samples corresponding to RR homozygotes, SR heterozygotes, SS homozygotes and negative controls (distilled water) were included in each assay. Since resistance to pyrethroids associated to amino acid substitutions in the VGSC is inherited as a recessive trait (Davies et al. 2007, González-Cabrera et al. 2013), mites carrying the mutant allele in homozygosis (RR) are considered as pyrethroid-resistant, whereas mites carrying the wild-type allele in homozygosis (SS) and the heterozygotes (SR) are considered as pyrethroid-susceptible mites.

## RESULTS

The samples (capped brood) used in this study were collected from apiaries located across the three provinces of the Comunitat Valenciana region (Spain). In the 2018 season, brood combs were collected from 90 apiaries (Table 2) while in the 2019 season they were collected from 89 apiaries (Table 3). As we need at least 120 live mites per sample to carry out the bioassays, not all the samples collected had sufficient mites. Hence, the level of parasitism required for bioassays was found in 58 % and 81 % of the samples in 2018 and 2019, respectively. On the other hand, since it is possible to carry out TaqMan® assays using a smaller number of mites collected either dead or alive, more analyses were carried out with this technique than with bioassays, resulting in 70 % and 94 % of the samples tested in 2018 and 2019 seasons, respectively.

**Table 2.**
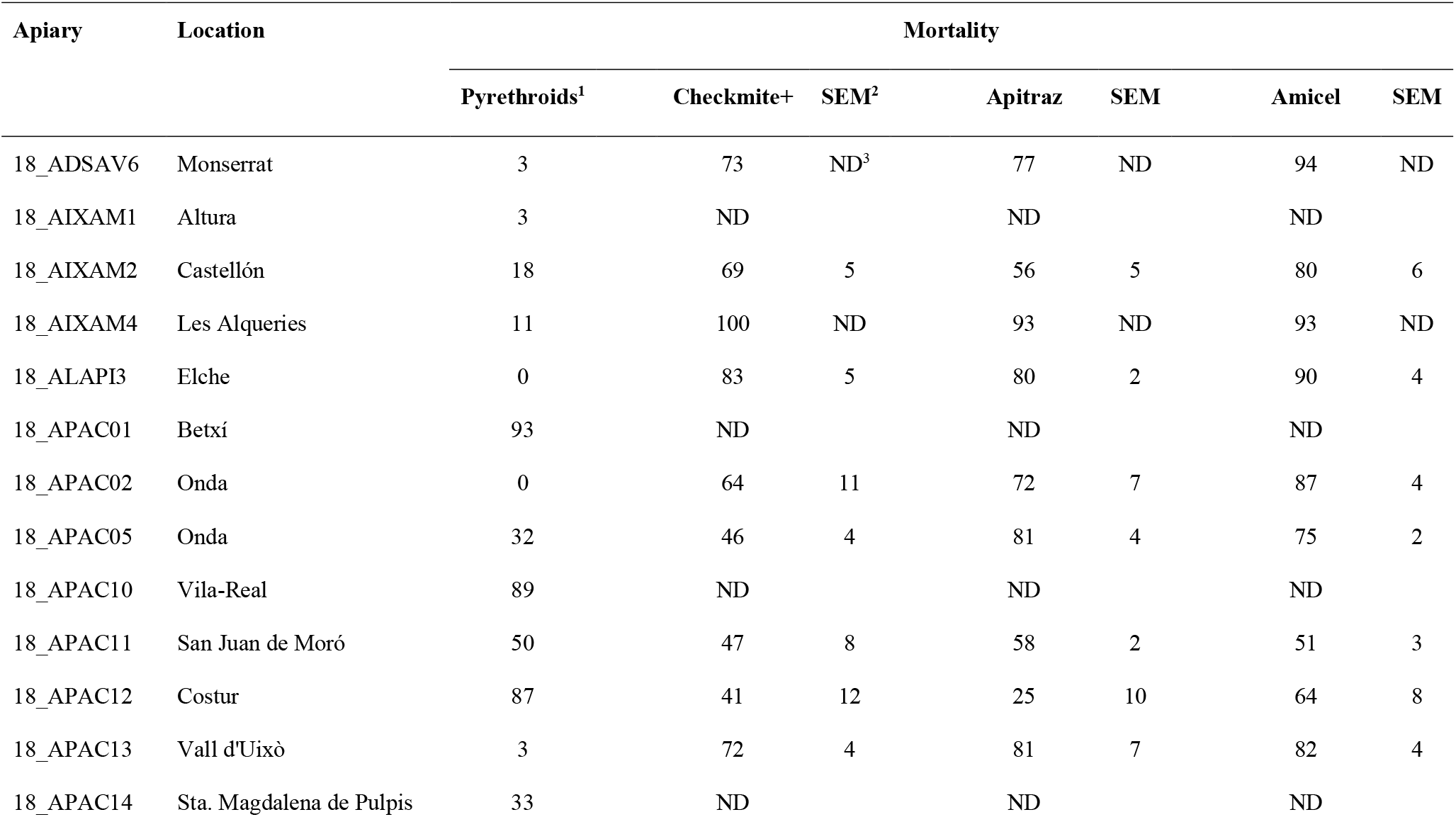
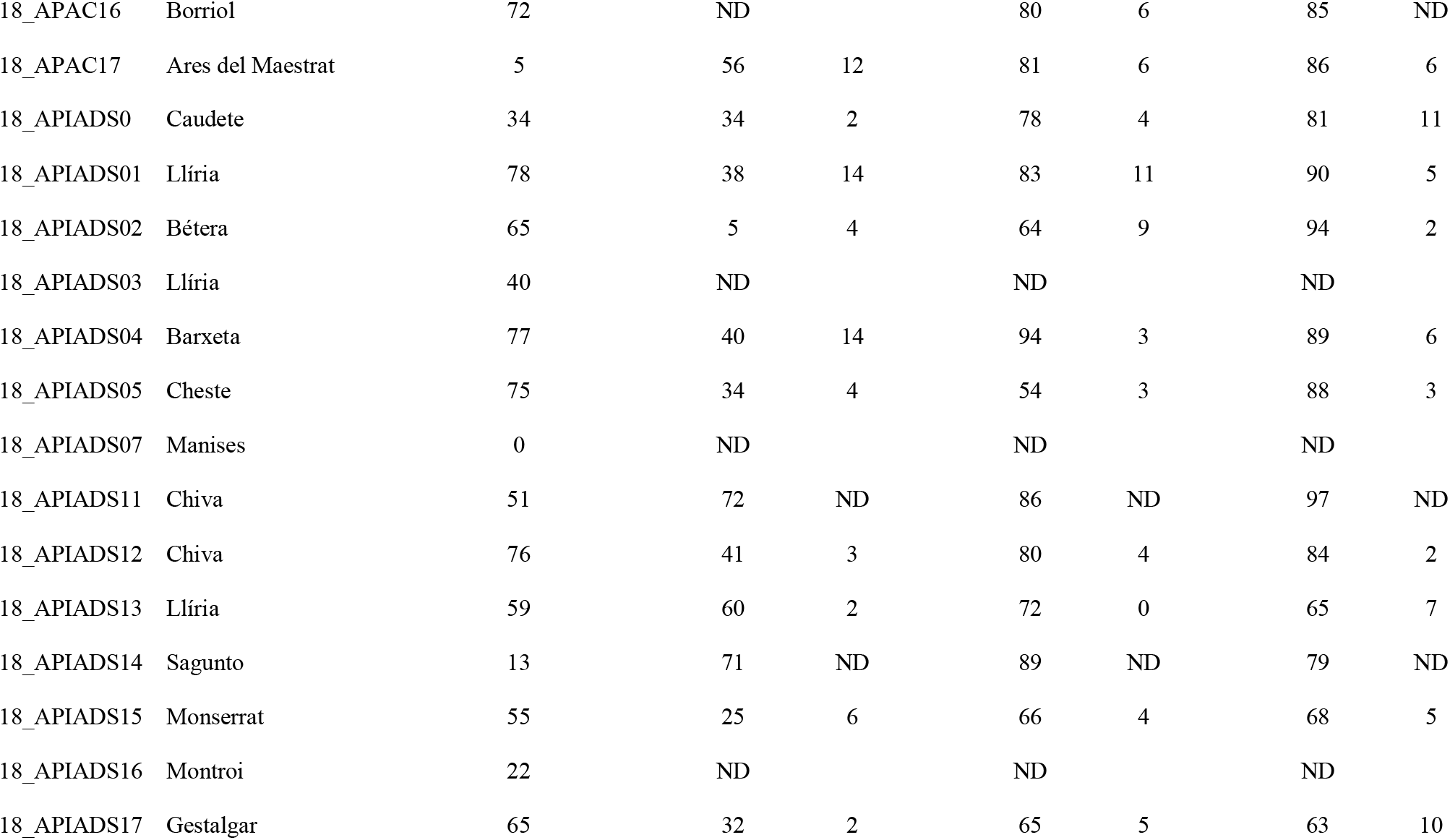
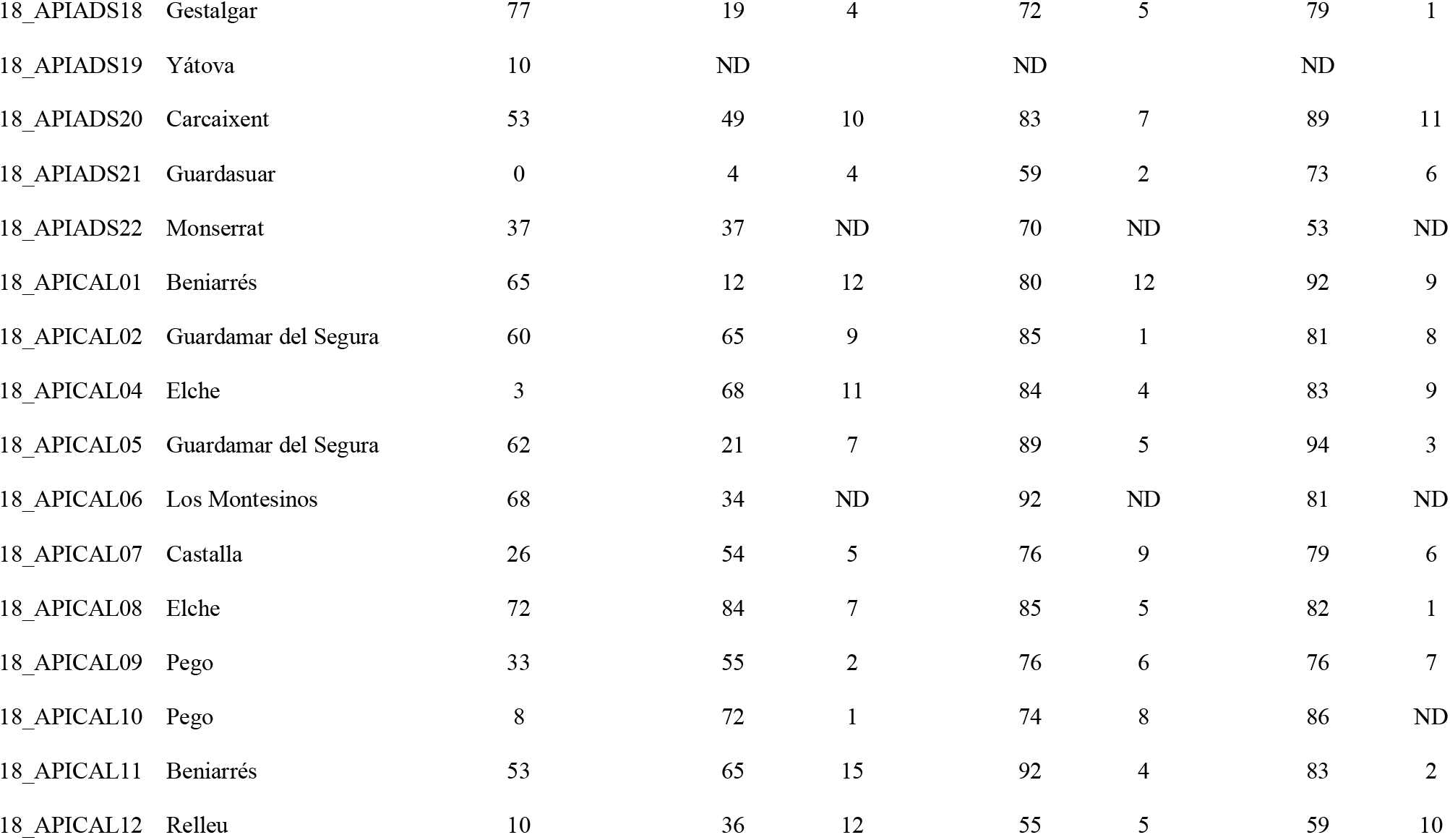
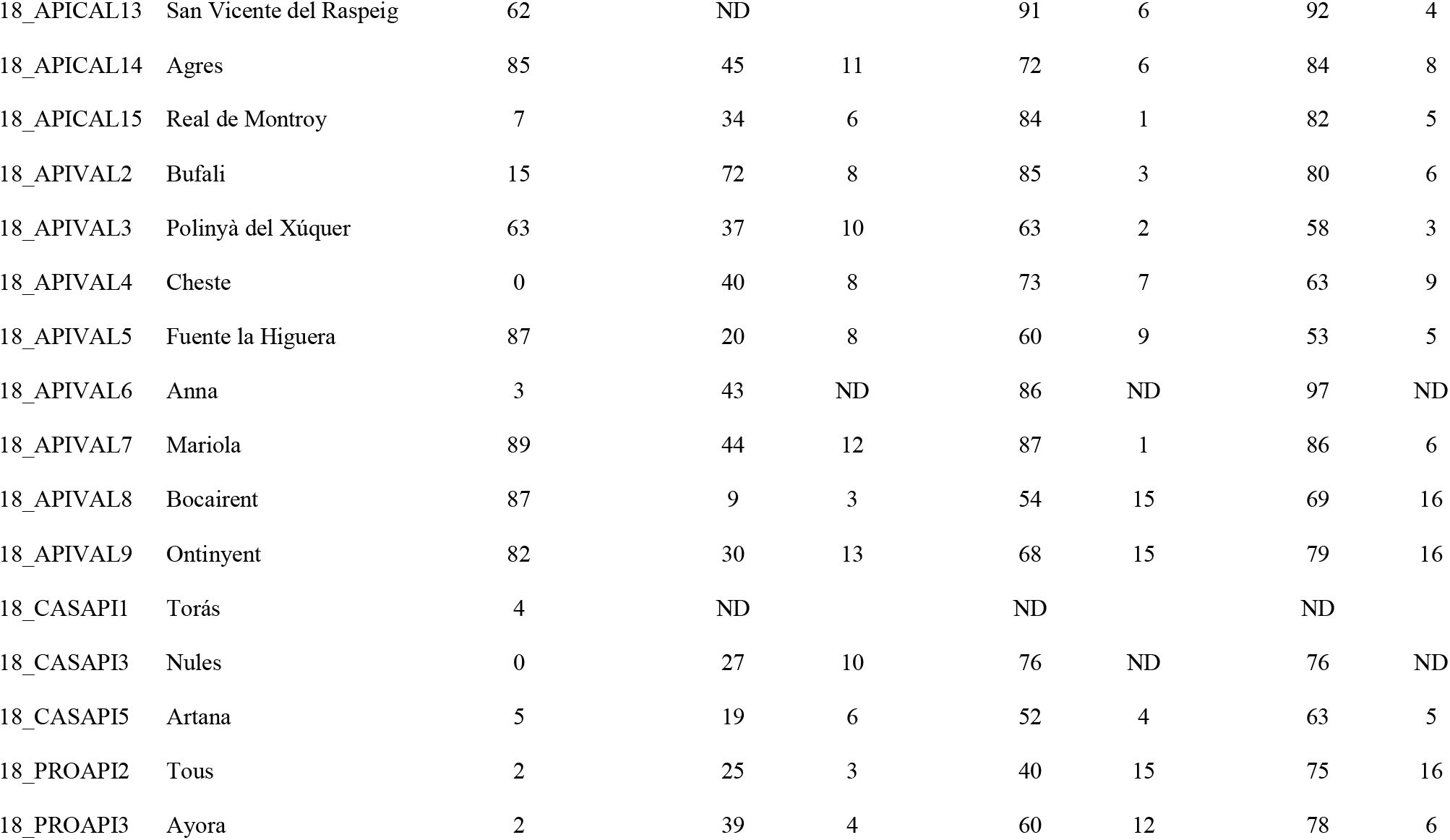
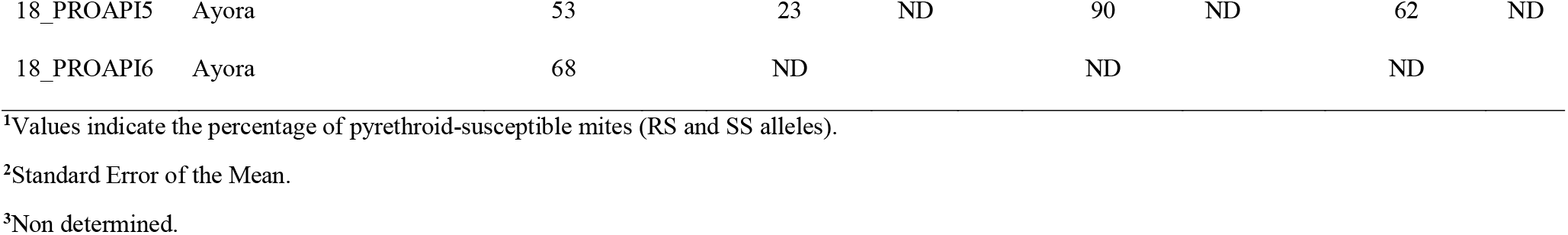
Sample locations and mortality of acaricides recorded in assays from 2018 season.

**Table 3.** Sample locations and mortality of acaricides recorded in assays from 2019 season.

| Apiary | Location | Mortality |  |  |  |  |  |  |
| --- | --- | --- | --- | --- | --- | --- | --- | --- |
|  |  | Pyrethroids<sup>1</sup> | Checkmite+ | SEM<sup>2</sup> | Apitraz | SEM | Amicel | SEM |
| 19_ADSAV01 | Llíria | 8 | ND<sup>3</sup> |  | ND |  | ND |  |
| 19_ADSAV02 | ND | 40 | 98 | ND | 83 | ND | 75 | ND |
| 19_AIXAM02 | Castellón | 0 | 71 | ND | 82 | ND | 86 | ND |
| 19_AIXAM04 | Altura | 6 | ND |  | ND |  | ND |  |
| 19_ALAPI01 | Guardamar del Segura | 12 | ND |  | ND |  | ND |  |
| 19_APAC01 | Torrechiva | 0 | ND |  | ND |  | ND |  |
| 19_APAC02 | Vistabella | 81 | ND |  | ND |  | ND |  |
| 19_APAC03 | Onda | 6 | 54 | 12 | 85 | 4 | 82 | 4 |
| 19_APAC04 | Alquerías del Niño Perdido | 10 | 64 | 2 | 72 | 5 | 83 | 7 |
| 19_APAC05 | Sant Joan de Moró | 24 | 67 | 6 | 75 | 5 | 76 | 2 |
| 19_APAC06 | Borriol | 54 | 73 | 1 | 90 | 3 | 84 | 1 |
| 19_APAC07 | Catí | 57 | 51 | 2 | 74 | 1 | 74 | 1 |
| 19_APAC08 | Xodos | 20 | 48 | 3 | 90 | 0 | 80 | 5 |
| 19_APAC09 | Cervera del Maestre | 35 | 44 | ND | 87 | ND | ND |  |
| 19_APAC10 | Costur | 27 | 40 | 13 | 82 | ND | 79 | ND |
| 19_APAC11 | Atzaneta del Maestrat | 20 | 63 | 5 | 80 | 0 | 80 | 0 |
| 19_APAC12 | Ares | 8 | 23 | 4 | 80 | 5 | 76 | 5 |
| 19_APAC13 | Torreblanca | 0 | 24 | 7 | 74 | 4 | 78 | 0 |
| 19_APIADS01 | Montroi | 10 | 72 | 4 | 65 | 14 | 84<sup>a</sup> | 2 |
| 19_APIADS02 | Casinos | 80 | 38 | 14 | 83 | 2 | 79 | 10 |
| 19_APIADS03 | Montserrat | 48 | 23 | ND | ND |  | ND |  |
| 19_APIADS04 | Higueruelas | 0 | ND |  | ND |  | ND |  |
| 19_APIADS05 | Picassent | 9 | ND |  | ND |  | ND |  |
| 19_APIADS06 | Montroi | 72 | 63 | 2 | 77 | 2 | 75 | 4 |
| 19_APIADS07 | Navarrés | 20 | 21 | ND | ND |  | ND |  |
| 19_APIADS08 | Valencia | 84 | 54 | 10 | 90 | 7 | 83 | 0 |
| 19_APIADS09 | Llíria | 90 | 23 | ND | ND |  | ND |  |
| 19_APIADS10 | Chiva | 54 | 50 | ND | 93 | ND | 82 | ND |
| 19_APIADS11 | Alzira | 57 | 63 | 8 | 89 | 2 | 76 | 8 |
| 19_APIADS12 | Llíria | 32 | 37 | 5 | 80 | 4 | 75 | 3 |
| 19_APIADS13 | Bunyol | 45 | 62 | 2 | 78 | 4 | 83 | 4 |
| 19_APIADS14 | Torrent | 60 | 82 | 2 | 86 | 1 | 80 | 1 |
| 19_APIADS15 | Chiva | 57 | 53 | ND | 82 | ND | ND |  |
| 19_APIADS16 | ND | 12 | ND |  | ND |  | ND |  |
| 19_APIADS17 | Alaquàs | 0 | 69 | 1 | 74 | 1 | 71 | 4 |
| 19_APIADS19 | Villarmarxant | 74 | 73 | 7 | 76 | 1 | 80 | ND |
| 19_APIADS20 | Ceste | 0 | 61 | 6 | 83 | 2 | 83 | 2 |
| 19_APIADS21 | Ceste | 0 | 60 | ND | 87 | ND | 80 | ND |
| 19_APIADS22 | Chiva | 47 | 52 | 3 | 82 | 0 | 83 | 1 |
| 19_APIADS23 | Chiva | 36 | 48 | ND | 83 | ND | 78 | ND |
| 19_APIADS24 | Bétera | 28 | 60 | 5 | 84 | 1 | 84 | 0 |
| 19_APIADS25 | Bétera | 32 | 55 | 4 | 73 | 1 | 75 | 1 |
| 19_APIADS26 | Bétera | 85 | 30 | 1 | 71 | 3 | 67 | 3 |
| 19_APIADS27 | Chiva | 77 | 56 | 7 | 81 | 4 | 78 | 3 |
| 19_APICAL01 | Molins | 18 | 67 | 11 | 85 | 4 | 90a | 3 |
| 19_APICAL02 | Rafol de Almunia | 3 | 66 | 2 | 68 | 4 | 74 | 2 |
| 19_APICAL03 | Gata de Gorjos | 78 | 33 | 5 | 66 | 6 | 62 | 13 |
| 19_APICAL04 | Benitachell | 47 | 44 | ND | 89 | ND | 93 | ND |
| 19_APICAL05 | Vilajoiosa | 95 | 52 | ND | 90 | ND | 83 | ND |
| 19_APICAL06 | San Miguel de Salinas | 11 | 40 | 3 | 84 | 5 | 87 | 1 |
| 19_APICAL07 | Torremanzanas | 92 | 46 | 7 | 86 | 1 | 80 | 1 |
| 19_APICAL08 | Torremanzanas | 0 | 59 | 6 | 80 | 3 | 77 | 3 |
| 19_APICAL09 | Finestrat | 0 | 53 | 1 | 82 | 4 | 82 | 1 |
| 19_APICAL10 | Agres | 85 | 69 | 1 | 85 | 3 | 69 | 1 |
| 19_APICAL11 | Castalla | 60 | 81 | 1 | 86 | 1 | 83 | 1 |
| 19_APICAL12 | Sella | 55 | 53 | ND | 64 | ND | 69 | ND |
| 19_APIVAL01 | Quesa | 0 | 60 | 4 | 72 | 4 | 77 | 1 |
| 19_APIVAL02 | La Font de la Figuera | 68 | 36 | 4 | 87 | 2 | 87 | 2 |
| 19_APIVAL03 | Favara | 5 | 31 | ND | 93 | ND | 93 | ND |
| 19_APIVAL04 | Carcaixent | 20 | 52 | 5 | 90 | 0 | 86 | 4 |
| 19_APIVAL05 | Sumacàrcer | 0 | 58 | 2 | 86 | 0 | 78 | 2 |
| 19_APIVAL06 | Bétera | 3 | 29 | 2 | 90 | 2 | 88 | 1 |
| 19_APIVAL07 | Albal | 36 | 57 | 3 | 89 | 1 | 83 | 4 |
| 19_APIVAL08 | Yátova | 2 | 67 | 4 | 83 | ND | 82 | ND |
| 19_APIVAL09 | Polinyà del Xúquer | 47 | ND |  | ND |  | ND |  |
| 19_APIVAL10 | Polinyà del Xúquer | 40 | 75 | 7 | 76 | 0 | 77 | 1 |
| 19_APIVAL11 | Bocairent | 28 | 34 | 3 | 76 | 1 | 77 | 0 |
| 19_APIVAL12 | Vall d'Alcalà | 5 | 68 | 2 | 76 | 5 | 65 | 6 |
| 19_APIVAL13 | Villena | 97 | 40 | ND | 83 | ND | 81 | ND |
| 19_CASAPI01 | Mas de Noguera | 49 | 23 | 4 | 79 | 3 | 69 | 4 |
| 19_CASAPI03 | Tales | 17 | 85 | 4 | 94 | ND | 72 | ND |
| 19_CASAPI04 | Sacañet | 20 | 75 | 2 | 72 | 2 | 75 | 2 |
| 19_PROAPI01 | Ayora | 5 | 67 | ND | 85 | ND | ND |  |
| 19_PROAPI02 | Énova | 0 | 38 | ND | 90 | ND | 83 | ND |
| 19_PROAPI03 | Orihuela | 97 | 68 | 1 | 76 | 0 | 72 | ND |
| 19_PROAPI04 | Requena | 89 | 54 | ND | 100 | ND | 72 | ND |
| 19_PROAPI05 | Moixent | 96 | 40 | 3 | 73 | 3 | 75 | 9 |
| 19_PROAPI06 | Requena | 2 | 85 | 6 | 85 | 2 | 77 | 2 |
| 19_PROAPI07 | Ayora | 0 | ND |  | ND |  | ND |  |
| 19_PROAPI08 | Ayora | 10 | 83 | ND | 87 | ND | 98 | ND |
| 19_PROAPI09 | Náquera | 65 | 61 | 1 | 74 | 1 | 89 | 0 |
| 19_PROAPI10 | Ayora | 72 | 69 | ND | 75 | ND | ND |  |
| 19_PROAPI11 | Ayora | 50 | ND |  | ND |  | ND |  |
| 19_PROAPI12 | Ayora | 32 | 61 | 10 | 84 | 3 | 79 | 2 |

<sup>1</sup>Values indicate the percentage of pyrethroid-susceptible mites (RS and SS alleles).
<sup>2</sup>Standard Error of the Mean.
<sup>3</sup>Non determined.

Results from bioassays conducted with Checkmite+ strips (coumaphos a.i.) showed some variability among samples, with a mean mortality of 50 % (± 21 SD) in 2018 (Fig. 2A), and 54 % (± 17 SD) in 2019 (Fig. 2B). Overall, the 75 % percentile value of approximately 66 % mortality obtained from the 122 apiaries tested in both seasons indicated that this product was less effective than expected according to the label.

**Figure 2.**
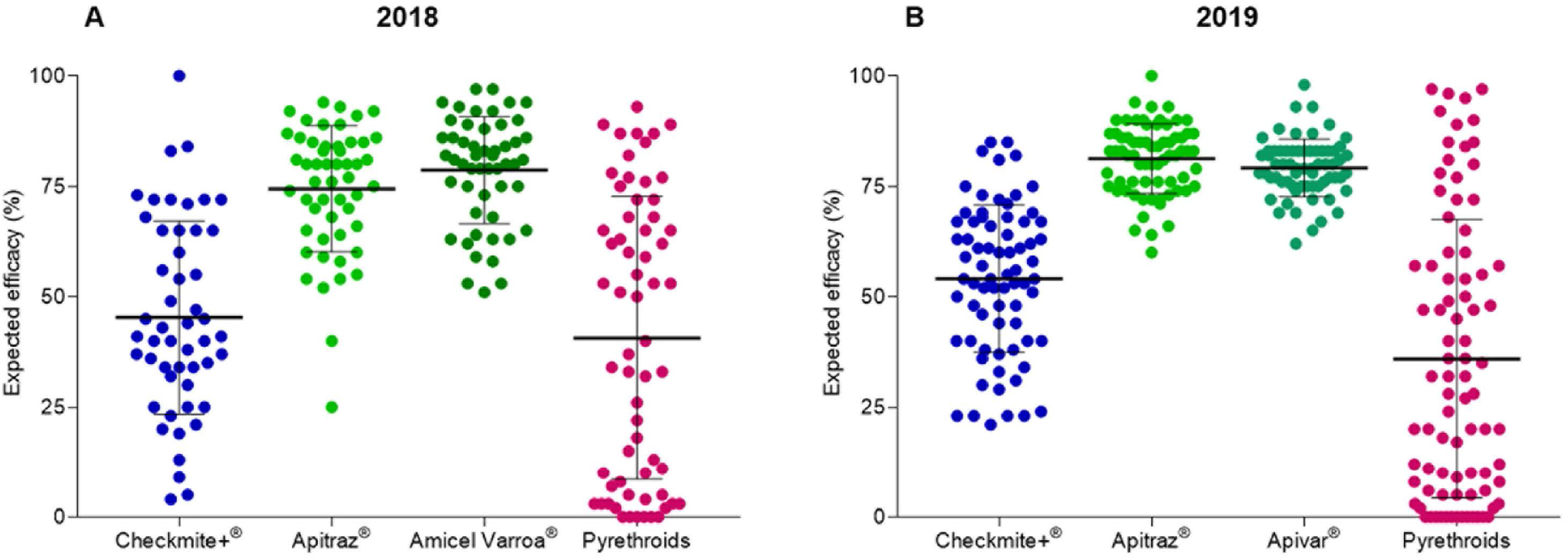
Expected efficacy of commercial acaricides against *Varroa destructor* in 2018 season (A) and 2019 season (B). For Checkmite+, Apitraz, Amicel and Apivar, the expected efficacy corresponds to mortality recorded in the bioassays. The expected efficacy of pyrethroids-based acaricides was estimated using the frequency of pyrethroid-resistant and susceptible mites after genotyping individual mites for the presence of different alleles of the mutation L925V at the *V. destructor* VGSC.

Three commercial acaricides based on amitraz were tested: Apitraz, Amicel Varroa and Apivar. Results from bioassays carried out with Apitraz in 2018 showed a mean mortality of 74 % (± 14 SD) (Fig. 2A). In that season, bioassays with Amicel Varroa showed a mean mortality of 79 % (± 12 SD) (Fig. 2A). Statistical analysis showed that these values were not significantly different (t test, *P* value > 0.05). In 2019, mean mortality with Apitraz and Apivar was 81 % (± 8 SD) and 79 % (± 7 SD), respectively (Fig. 2B), again with no significant differences between them (t test, *P* value > 0.05). Therefore, according to our data, the mortality of the three amitraz-based acaricides tested was found to be similar across the study.

The expected efficacy of pyrethroids based acaricides against *V. destructor* was estimated using a TaqMan® genotyping assay. The frequency of pyrethroid-resistant and susceptible mites was determined for each sample after genotyping 40 individual mites for the presence of different alleles of the mutation L925V at the *V. destructor* VGSC. When TaqMan® assays were conducted, a wide range of allele frequency patterns was found in the apiaries evaluated in both, 2018 and 2019 seasons. The estimated mean efficacy of pyrethroids was 41 % (± 32 SD) in samples from 2018 (Fig. 2A), and 36 % (± 32 SD) in samples from 2019 (Fig. 2B). The Standard Deviation of these means corroborated the high dispersion of pyrethroid expected efficacies throughout the region, ranging from zero (in apiaries with all mites resistant to pyrethroids), to 97 % in some apiaries with almost all mites susceptible to the acaricide.

The efficacy of the different acaricides in each apiary was weighed according to its geographic location (Fig. 3 and 4). Our data show that there is no geographic dependent pattern of expected efficacies to the acaricides tested, as it can be noticed looking at the different mortality values estimated in apiaries at nearby locations (Fig. 3 and 4; Tables 2 and 3).

**Figure 3.**
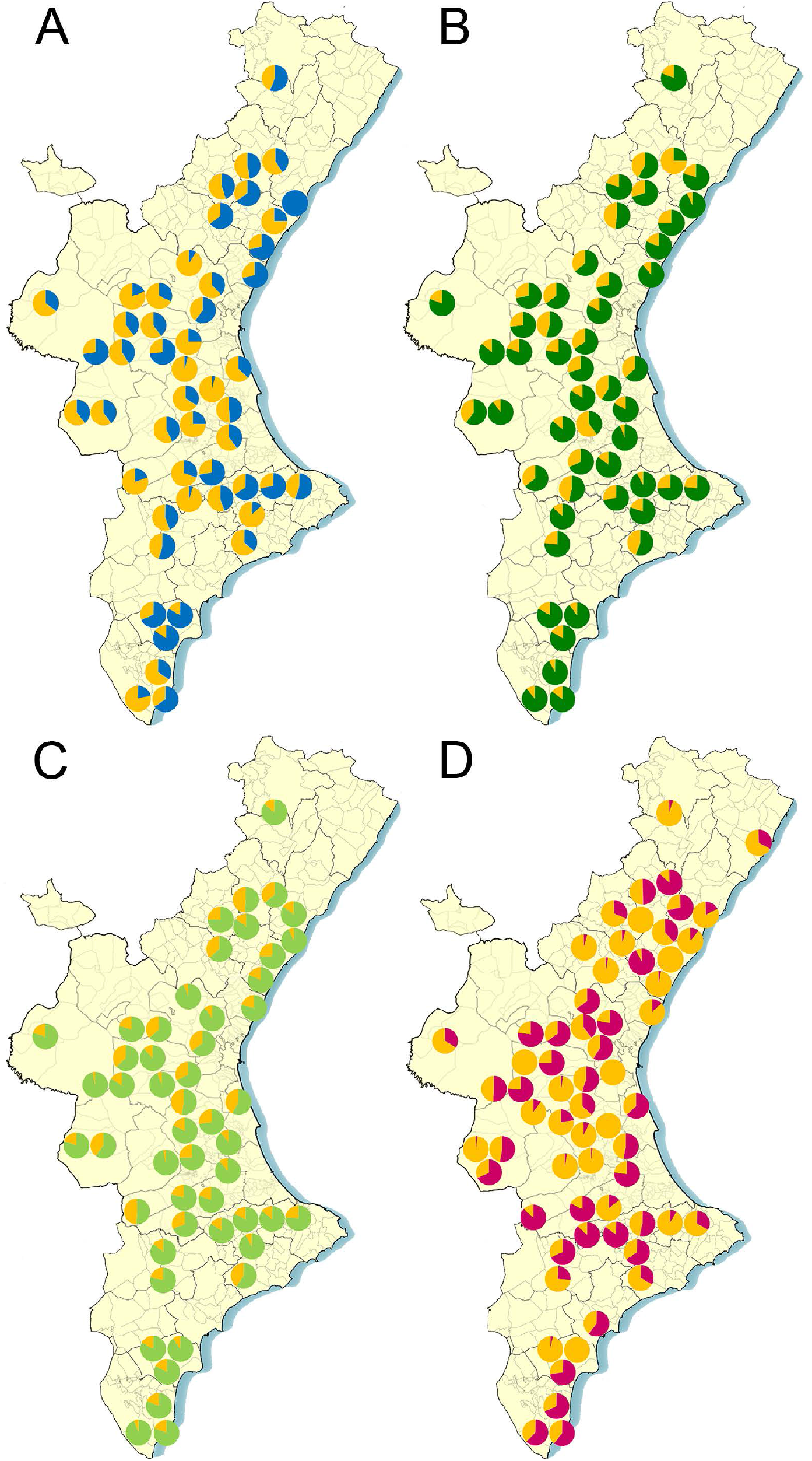
Sampling locations and expected efficacy (expressed as percentage) of commercial acaricides against *Varroa destructor* in 2018 season. A) Coumaphos (efficacy in blue); B) Apitraz (efficacy in dark green); C) Amicel Varroa (efficacy in light green); D) Pyrethroids (efficacy in pink).

**Figure 4.**
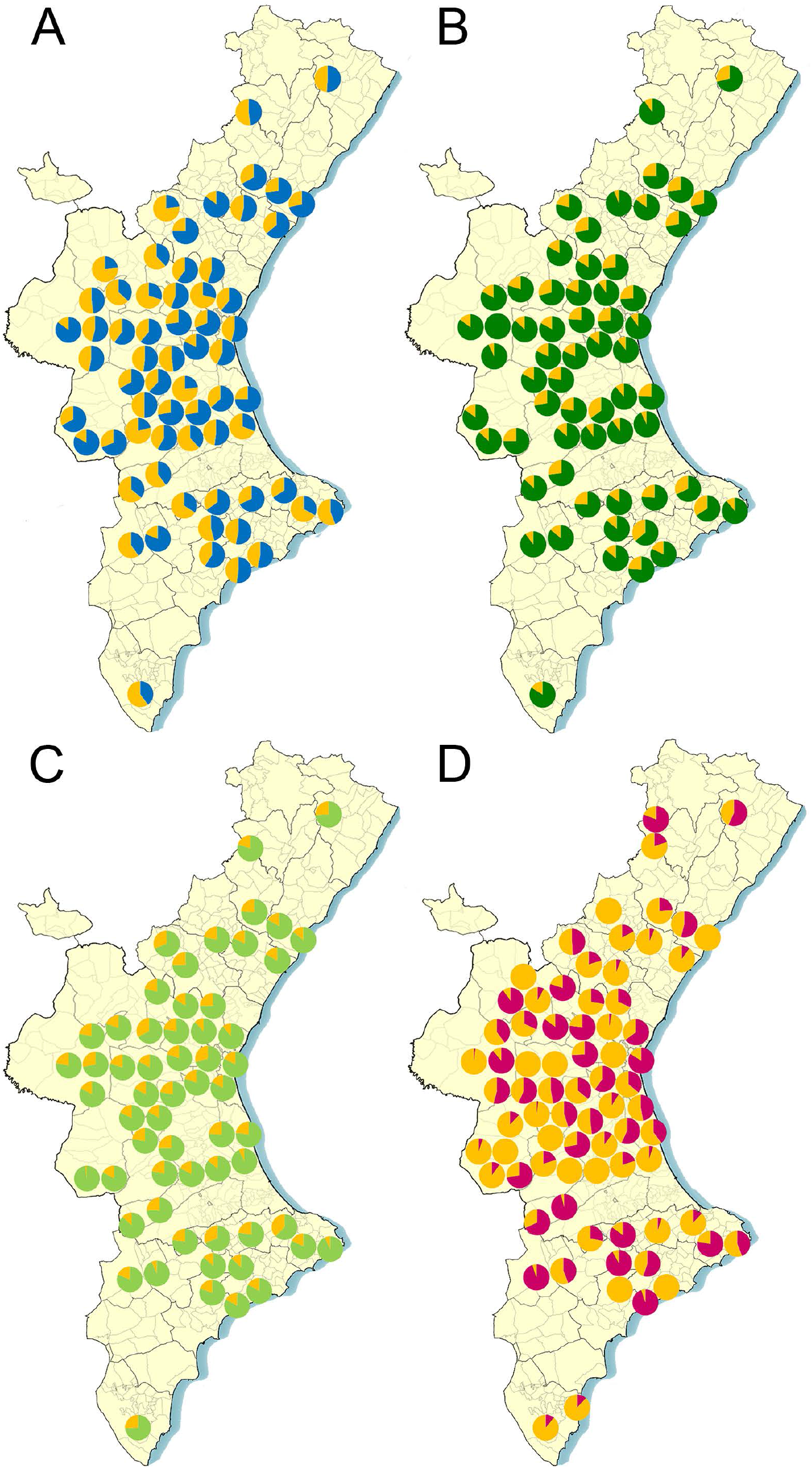
Sampling locations and expected efficacy (expressed as percentage) of commercial acaricides against *Varroa destructor* in 2019 season. A) Coumaphos (efficacy in blue); B) Apitraz (efficacy in dark green); C) Apivar (efficacy in light green); D) Pyrethroids (efficacy in pink).

Along with the brood combs, information about treatment history was collected from each apiary. These data showed that most of the beekeeping operations used amitraz-based acaricides (88 % of the treatments), while the use of other treatment regimens such as those based on pyrethroids and soft acaricides was much lower, representing 5 % and 7 % of the total treatments, respectively. (Fig. 5).

**Figure 5.**
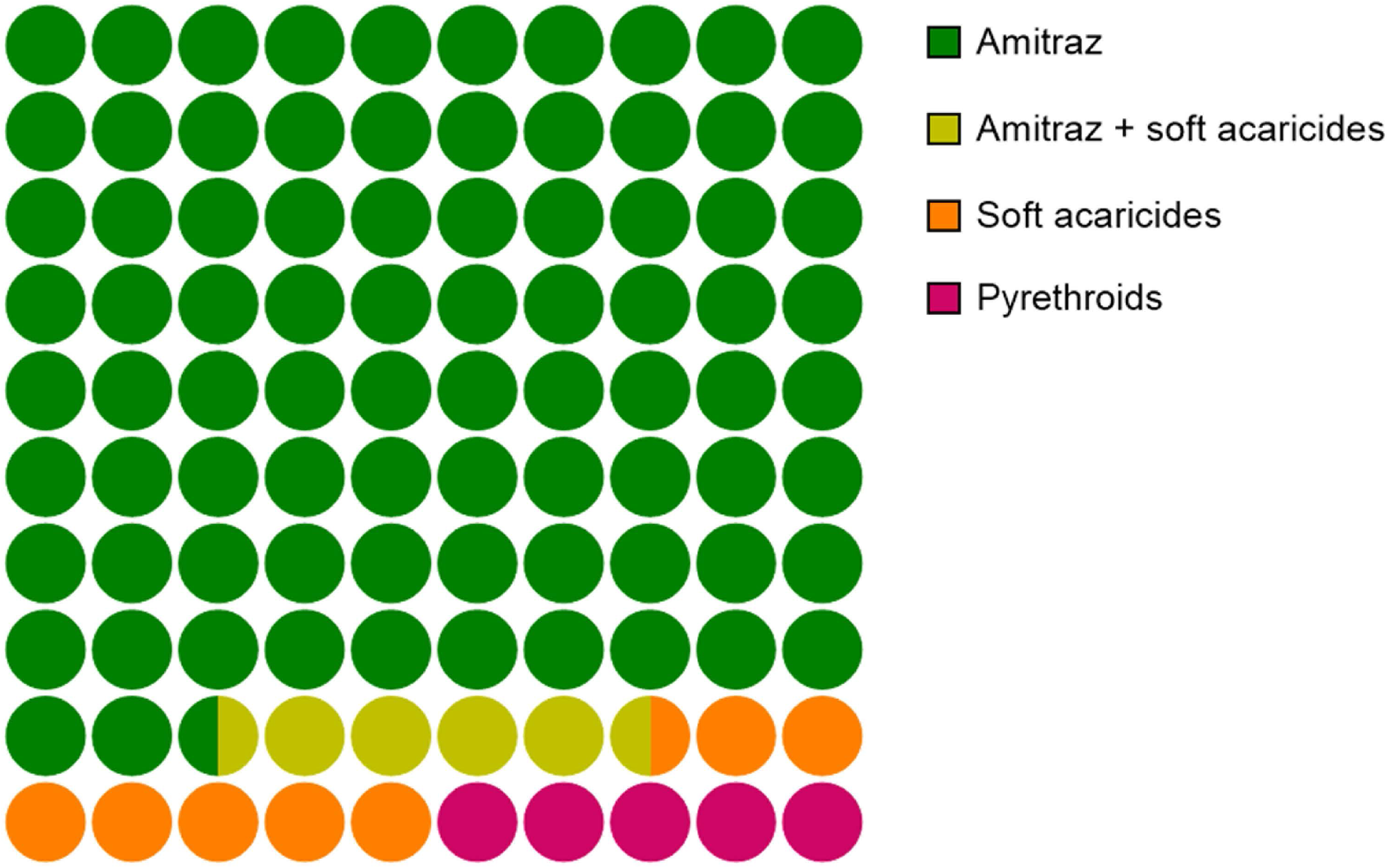
Acaricide treatments (%) in the apiaries that provided samples for this study in 2018 and 2019 seasons.

## DISCUSSION

The data presented here is the first comprehensive and large-scale monitoring study describing the situation of the resistance to acaricides in populations of *V. destructor*. The analyzed apiaries belong to the Comunitat Valenciana, one of the most prominent apicultural regions in Spain. Moreover, since migratory beekeeping is a common practice among beekeepers, it is possible to hypothesize that the current situation in this region may resembled that of the rest of the country.

Varroosis is one of the major threats to the viability of apiculture, not only in Spain, but worldwide. Beekeepers are currently struggling to find good alternatives to reach an adequate management of the mite since there are very few active ingredients and formulations authorized to battle the disease. Therefore, it is crucial that the acaricides already in the market remain effective for as long as possible, although the evolution of resistance is currently jeopardizing this aim. If resistance to an acaricide is detected and the treatments with this compound are not discontinued immediately, resistant individuals will be selected and they will become more frequent in the population, leading to therapeutic failures in more and more places as resistance spreads. For this reason, programs to monitor the efficacy of acaricide products are of critical importance.

The huge volume of apiaries in Spain require an adequate management of the active ingredients for *V. destructor* control, so that the most appropriated for each operation is used depending on the susceptibility of the mite population to each acaricide. However, there is no official program in this country recording the efficacy of treatments, nor the monitoring of possible outbreaks of resistance to these treatments. In a recent study, bioassays with Checkmite, Apivar and Apistan were conducted with samples collected in seven Spanish locations (Higes et al. 2020), providing a general idea of the acaricide efficacy in these apiaries. Aiming to perform a more thorough study, we carried out analyses in two consecutive years with a significantly higher number of samples, covering almost completely the area dedicated to beekeeping in this region. To obtain data from a large number of apiaries, we coordinate with the government of the Comunitat Valenciana region (Generalitat Valenciana) and nine sanitary defence groups (ADS) of the beekeeping sector. As a consortium, we started a program to evaluate the efficacy of the authorized active ingredients to control *V. destructor* in apiaries of this Spanish region. The study was carried out during 2018 and 2019 beekeeping seasons testing the three groups of hard acaricides authorized in Spain: coumaphos, amitraz and pyrethroids.

Coumaphos has been widely used for many years as an active ingredient in Checkmite+ commercial strips. In Spain, a reduction of this product efficacy to control *V. destructor* parasitism was detected a few years ago, with mean efficacies of about 70 to 80 % (Sánchez Escudero and Fernández Tejedor 2016, Calatayud et al. 2018). Actually, the product’s distributor in Spain, Bayer Hispania, S.L. (Animal Health), issued a statement informing beekeepers of a possible lack of Varroa sensitivity to coumaphos based on preliminary results obtained in a study with this acaricide in three areas of central and northern Spain (https://www.aeapicultores.org/wp-content/uploads/2017/03/Comunicado_Bayer_Apicultores.pdf. From then on, the use of this acaricide declined dramatically and even the Spanish Association of Veterinary Specialists in Health and Bee Production (AVESPA acronym) demanded the withdrawal of the product from the market and discouraged its use among beekeepers (http://www.colvet.es/node/2663). Tracking the efficacy of Checkmite+ strips with mites from the apiaries of this study confirmed the reduction of its efficacy to control Varroa. According to our data, the mortality observed with this acaricide varied considerably from one apiary to another (Fig. 3 and 4; Tables 2 and 3), although in all cases the expected efficacy would be lower than that indicated by the manufacturer. The significant variation in the mortality registered for coumaphos in the bioassays throughout the study seems to indicate that the reduced efficacy is due to the presence of mites resistant to this acaricide in the hives. Actually, our previous study with mites sampled in different Spanish regions also found that resistance to coumaphos is evolving in this country (Higes et al. 2020). *Varroa destructor* populations resistant to coumaphos were previously reported in America (Elzen and Westervelt 2002, Maggi et al. 2009, Maggi et al. 2011). In particular, the resistance reported in one of these studies does not seem reversible after stopping the treatments with coumaphos (Mitton et al. 2018). Our results support this hypothesis since the apiaries in this study had not been treated with coumaphos for several years and still part of the populations remained insensitive to this compound. The accumulation and persistence of coumaphos residues in beeswax for a long time (Calatayud-Vernich et al. 2018), could favour a constant selection pressure in the colonies, reducing the possibility of reversing the resistance. However, to rule out this hypothesis it may be necessary to identify the molecular mechanism of the resistance, which may also be of help to design molecular tools to identify coumaphos-resistant individuals in the populations.

On the other hand, the mechanism of *V. destructor* resistance to pyrethroids is well described, so a TaqMan® allelic discrimination assay was used to identify the resistant mites carrying a mutation in the 925 position of the VGSC as described by González-Cabrera et al. (2013). We used the proportion of susceptible and pyrethroid-resistant individuals in each apiary to estimate the efficacy that a pyrethroid treatment would have. The results also showed great diversity in the proportion of resistant mites among apiaries. Remarkably, bees on many operations were parasitized only by pyrethroid-resistant mites, but there were also others with most of the mites labelled as susceptible (Fig. 3 and 4; Tables 2 and 3). The long-track record of treatments only with pyrethroid-based acaricides and their accumulation in beeswax (Calatayud-Vernich et al. 2018) is a likely explanation for the evolution of resistance. Although it has been suggested several times that the mutations associated with the resistance to pyrethroids in Varroa cause a reduced fitness in the mites (Milani and Della Vedova 2002, González-Cabrera et al. 2016, González-Cabrera et al. 2018), it seems that this is not enough to remove completely the resistant mites. The Varroa populations may contain a remnant of resistant individuals that would be quickly selected as soon as the treatment with pyrethroids are reinstated. Our data support this hypothesis since the mites collected from apiaries reporting the last treatment with pyrethroids were mainly labelled as resistant (data not shown). This observation is again confirming the idea that, although it is advisable to use pyrethroids to control Varroa in an IPM context when the frequency of resistant individuals is sufficiently low, it is very important to avoid applying continuous treatments with pyrethroid-based acaricides.

In the case of both, coumaphos and pyrethroids, no geographical distribution associated with the resistance was observed. On the contrary, the estimated efficacy for these treatments seemed to be randomly distributed. This might be surprising at first sight, but it can be explained by the idiosyncrasy of beekeeping in this region, which beekeepers mainly keeping hives that are seasonally moved across the country. The movement of hives to different areas is most certainly involving a transfer of parasitized bees from the migrant hives to those already in the transient settlement and vice versa. This can justify that each apiary has a specific pattern of acaricide efficacy depending on both, the treatments applied and the places they have traveled.

Apitraz, Apivar and Amicel Varroa, commercial products containing amitraz as an active ingredient, are widely used in Spain. The assays conducted showed that the expected efficacy in the field would be very similar for the three of them, and it would also be the highest amongst the different active ingredients tested in this study (Fig. 3 and 4; Tables 2 and 3). The low variation recorded after assaying amitraz in multiple apiaries is an indication of its consistency as acaricide across the country. However, given that our data indicate that the expected efficacy would be below 90 % in most cases, it is clear that the products are performing below the expected efficacy according to the label. These data is in agreement with the actual efficacy of amitraz-based product recorded by beekeepers in the field, with values ranging from 60 to 96 % (Calatayud et al. 2018). Hence, it is possible to anticipate an evolution of resistance to amitraz in the coming seasons unless there is a significant change in the management strategies, currently based on the intensive use of mainly amitraz many times per year. Repeated use of a single acaricide exerts an enormous selection pressure on the mites. If resistance to amitraz evolves, it will be much more difficult to control the parasite, since the other commercial products will not be effective in all apiaries. The recommendations to delay the evolution of resistance encourage the rotation of products with different mode of action (IRAC; https://www.irac-online.org/). Therefore, a more rational use of current acaricides would be desirable, alternating the application of different active ingredients in consecutive seasons. To decide whether a given treatment would be successful in each apiary, it is crucial to monitor the efficacy of the acaricidal compounds in each of them. In this way, the effective products could be rotated, avoiding selection pressures with treatments that can lead to an increase in the frequency of resistant mites.

The transfer from Academia to the field is one of the priorities of this work. The final objective was to provide the beekeeping sector with the information obtained in this study. The results from this project were disclosed in informative talks to groups of beekeepers. Moreover, tailored reports with the expected efficacy of acaricide treatments in each apiary were sent to the professionals in charge of the different ADS for them to discuss with the relevant beekeeper the best management approach for controlling the mites in their apiaries, considering also the previous history of treatments.

Initiatives such as the Honey Bee Health Coalition (https://honeybeehealthcoalition.org/) and the Bee informed partnership (https://beeinformed.org/) in the USA (coalitions of researchers, advisors, and stakeholders from various sides of the honey bee related industry) are good examples of associations that encourage the flow of information among the different actors in the beekeeping world. The rational use of pesticides to manage *V. destructor* is a joint responsibility of public institutions, industry, Academia and the beekeeping sector. Only by acting in a coordinated manner the efficacy of treatments may be prolonged and the evolution of resistances that threaten the viability of apiculture may be delayed.

## ACKNOWLEDGMENTS

The authors want to thank all beekeepers and beekeeper associations for contributing samples for this study and to the Comunitat Valenciana Regional Government (Agriculture and Livestock Subdivision) for helping with the coordination of beekeeper associations and sample collection.

## Notes

**FUNDING**: Joel González-Cabrera was supported by the Spanish Ministry of Economy and Competitiveness, Ramón y Cajal Program (grant: RYC-2013-261 13834). The work at the Universitat de València was funded by the Spanish Ministry of Economy and Competitiveness (grant: CGL2015-65025-R, MINECO/FEDER, UE), the Spanish Ministry of Agriculture, Fisheries and Food (grants: 20180020000920 and 201900700000555), the Spanish Ministry of Science, Innovation and Universities (grant: RTI2018-095120-B-100) and Conselleria de Agricultura, Desarrollo Rural, Emergencia Climática y Transición Ecológica - Agriculture and Livestock Subdivision (grants: CNME18/71480/12 and CNME19/71480/46)

### Competing Interest Statement

The authors have declared no competing interest.

### Summary of Updates

This version has been updated to include one author and to update funding information

## REFERENCES

Boecking, O., and E. Genersch. 2008. Varroosis - the ongoing crisis in bee keeping. Journal Fur Verbraucherschutz Und Lebensmittelsicherheit-Journal of Consumer Protection and Food Safety 3: 221–228.

Calatayud-Vernich, P., F. Calatayud, E. Simó, and Y. Picó. 2018. Pesticide residues in honey bees, pollen and beeswax: Assessing beehive exposure. Environmental Pollution 241: 106–114.

Calatayud, F., E. Simó, and P. Domingo. 2018. Hacia un control integrado y sostenible de Varroa (1). Apicultura Ibérica 27: 15–25.

Davies, T. G. E., L. M. Field, P. N. R. Usherwood, and M. S. Williamson. 2007. DDT, pyrethrins, pyrethroids and insect sodium channels. IUBMB life 59: 151–162.

Elzen, P. J., and D. Westervelt. 2002. Detection of coumaphos resistance in *Varroa destructor* in Florida. Am Bee J 142: 291–292.

Elzen, P. J., F. A. Eischen, J. B. Baxter, J. Pettis, G. W. Elzen, and W. T. Wilson. 1998. Fluvalinate resistance in *Varroa jacobsoni* from several geographic locations. Am Bee J 138: 674–676.

Genersch, E., W. von der Ohe, H. Kaatz, A. Schroeder, C. Otten, R. Büchler, S. Berg, W. Ritter, W. Mühlen, S. Gisder, M. Meixner, G. Liebig, and P. Rosenkranz. 2010. The German bee monitoring project: a long term study to understand periodically high winter losses of honey bee colonies. Apidologie 41: 332–352.

González-Cabrera, J., T. G. E. Davies, L. M. Field, P. J. Kennedy, and M. S. Williamson. 2013. An amino acid substitution (L925V) associated with resistance to pyrethroids in *Varroa destructor*. Plos One 8: e82941.

González-Cabrera, J., S. Rodríguez-Vargas, T. G. E. Davies, L. M. Field, D. Schmehl, J. D. Ellis, K. Krieger, and M. S. Williamson. 2016. Novel Mutations in the voltage-gated sodium channel of pyrethroid-resistant *Varroa destructor* populations from the Southeastern USA. Plos One 11: e0155332.

González-Cabrera, J., H. Bumann, S. Rodríguez-Vargas, P. J. Kennedy, K. Krieger, G. Altreuther, A. Hertel, G. Hertlein, R. Nauen, and M. S. Williamson. 2018. A single mutation is driving resistance to pyrethroids in European populations of the parasitic mite, *Varroa destructor*. Journal of Pest Science 91: 1137–1144.

Gracia-Salinas, M. J., M. Ferrer-Dufol, E. Latorre-Castro, C. Monero-Manera, J. A. Castillo-Hernández, J. Lucientes-Curd, and M. A. Peribanez-López. 2006. Detection of fluvalinate resistance in *Varroa destructor* in Spanish apiaries. J Apicult Res 45: 101–105.

Guzmán-Novoa, E., L. Eccles, Y. Calvete, J. Mcgowan, P. G. Kelly, and A. Correa-Benítez. 2010. *Varroa destructor* is the main culprit for the death and reduced populations of overwintered honey bee (*Apis mellifera*) colonies in Ontario, Canada. Apidologie 41: 443–450.

Higes, M., R. Martín-Hernández, C. S. Hernández-Rodríguez, and J. González-Cabrera. 2020. Assessing the resistance to acaricides in *Varroa destructor* from several Spanish locations. Parasitol Res.

Hubert, J., M. Nesvorna, M. Kamler, J. Kopecky, J. Tyl, D. Titera, and J. Stara. 2014. Point mutations in the sodium channel gene conferring tau-fluvalinate resistance in *Varroa destructor*. Pest Manag Sci 70: 889–894.

Kamler, M., M. Nesvorna, J. Stara, T. Erban, and J. Hubert. 2016. Comparison of tau-fluvalinate, acrinathrin, and amitraz effects on susceptible and resistant populations of *Varroa destructor* in a vial test. Exp Appl Acarol 69: 1–9.

Llorente, J. 2003. Principales enfermedades de las abejas, Ministerio de Agricultura, Pesca y Alimentación.

Maggi, M. D., S. R. Ruffinengo, P. Negri, and M. J. Eguaras. 2010. Resistance phenomena to amitraz from populations of the ectoparasitic mite *Varroa destructor* of Argentina. Parasitol Res 107: 1189–1192.

Maggi, M. D., S. R. Ruffinengo, N. Damiani, N. H. Sardella, and M. J. Eguaras. 2009. First detection of *Varroa destructor* resistance to coumaphos in Argentina. Experimental and Applied Acarology 47: 317–320.

Maggi, M. D., S. R. Ruffinengo, Y. Mendoza, P. Ojeda, G. Ramallo, I. Floris, and M. J. Eguaras. 2011. Susceptibility of *Varroa destructor* (Acari: Varroidae) to synthetic acaricides in Uruguay: Varroa mites’ potential to develop acaricide resistance. Parasitol Res 108: 815–821.

Milani, N. 1995. The resistance of *Varroa-Jacobsoni* Oud to pyrethroids - a laboratory assay. Apidologie 26: 415–429.

Milani, N., and G. Della Vedova. 2002. Decline in the proportion of mites resistant to fluvalinate in a population of *Varroa destructor* not treated with pyrethroids. Apidologie 33: 417–422.

Mitton, G. A., N. Szawarski, F. Ramos, S. Fuselli, F. R. Meroi Arcerito, M. J. Eguaras, S. R. Ruffinengo, and M. D. Maggi. 2018. *Varroa destructor*: when reversion to coumaphos resistance does not happen. J Apicult Res 57: 536–540.

Muñoz, I., E. Garrido-Bailón, R. Martín-Hernández, A. Meana, M. Higes, and P. De la Rúa. 2015. Genetic profile of Varroa destructor infesting Apis mellifera iberiensis colonies. J Apicult Res 47: 310–313.

Ramsey, S. D., R. Ochoa, G. Bauchan, C. Gulbronson, J. D. Mowery, A. Cohen, D. Lim, J. Joklik, J. M. Cicero, J. D. Ellis, D. Hawthorne, and D. vanEngelsdorp. 2019. *Varroa destructor* feeds primarily on honey bee fat body tissue and not hemolymph. Proc Natl Acad Sci U S A 116: 1792–1801.

Rinkevich, F. D. 2020. Detection of amitraz resistance and reduced treatment efficacy in the Varroa Mite, *Varroa destructor*, within commercial beekeeping operations. Plos One 15:e0227264.

Rodríguez-Dehaibes, S. R., G. Otero-Colina, V. P. Sedas, and J. A. V. Jiménez. 2005. Resistance to amitraz and flumethrin in *Varroa destructor* populations from Veracruz, Mexico. J Apicult Res 44: 124–125.

Rosenkranz, P., P. Aumeier, and B. Ziegelmann. 2010. Biology and control of *Varroa destructor*. J Invertebr Pathol 103: S96–S119.

Sammataro, D., P. Untalan, F. Guerrero, and J. Finley. 2005. The resistance of varroa mites (Acari : Varroidae) to acaricides and the presence of esterase. Int J Acarol 31: 67–74.

Sánchez Escudero, M. D., and T. Fernández Tejedor. 2016. Lucha frente a la varroosis en colmenas Layens.

Solignac, M., J. M. Cornuet, D. Vautrin, Y. Le Conte, D. Anderson, J. Evans, S. Cros-Arteil, and M. Navajas. 2005. The invasive Korea and Japan types of *Varroa destructor*, ectoparasitic mites of the Western honeybee (*Apis mellifera*), are two partly isolated clones. Proceedings of the Royal Society B-Biological Sciences 272: 411–419.

Underwood, R., and M. López-Uribe. 2020. Methods to control Varroa mites: an Integrated Pest Management approach. Penn State University.

